# Botulinum-neurotoxin-like sequences identified from an *Enterococcus* sp. genome assembly

**DOI:** 10.1101/228098

**Authors:** Charles H.D. Williamson, Theresa J. Smith, Brian T. Foley, Karen Hill, Paul Keim, Jason W. Sahl

## Abstract

Botulinum neurotoxins (BoNTs) are produced by diverse members of the *Clostridia* and result in a flaccid paralysis known as botulism. Exploring the diversity of BoNTs is important for the development of therapeutics and antitoxins. Here we describe a novel, *bont*-like gene cluster identified in a draft genome assembly for *Enterococcus* sp. 3G1_DIV0629 by querying publicly available genomic databases. The *bont*-like gene is found in a gene cluster similar to known *bont* gene clusters. Protease and binding motifs conserved in known BoNT proteins are present in the newly identified BoNT-like protein; however, it is currently unknown if the BoNT-like protein described here is capable of targeting neuronal cells resulting in botulism.

## Introduction

Diverse members of the genus *Clostridium* produce botulinum neurotoxins (BoNTs), which are some of the most toxic substances currently known. These neurotoxins cause botulism, a flaccid paralysis affecting mammals and birds, and are classified as Select Agents(https://www.selectagents.gov/SelectAgentsandToxinsList.html). BoNTs are also used for therapeutic purposes to treat a variety of conditions (*1*-*5*). Known BoNTs have been classified by antigenic cross reactivity, resulting in seven serotypes (types A-G). Within some serotypes, extensive diversity has been identified; toxin subtypes have been described based upon amino acid sequence composition (*6*). Researchers have worked to understand the diversity of BoNTs in order to develop therapeutics and to ensure that appropriate antitoxins are available. Recently, novel BoNT-like sequences have been identified using bioinformatics tools to search ever-increasing whole genome sequence data (*7*-*9*).

Botulinum neurotoxin genes (*bont*) are clearly mobile and can be located on the chromosome, on plasmids, or on bacteriophage (*10*, *11*). The composition of the toxin-gene clusters includes a number of accessory genes. A nontoxin/nonhemagglutinin gene (*ntnh*), which encodes a protein that protects the BoNT from degradation in the intestine (*12*), is directly adjacent to the botulinum neurotoxin gene in all known botulinum neurotoxin gene clusters. Other accessory genes are the genes encoding hemagglutinin proteins (*ha*+ clusters) that play a role in the BoNT crossing the intestinal barrier into the blood stream (*13*, *14*) or the open reading frames (*orfX*+ clusters) that are of unknown function. Additional genes that may be present in botulinum toxin gene clusters include a *botR* gene, a *lycA* gene and/or a *p47* gene. Interestingly, a *bont*-like gene (labelled BoNT/Wo by Zornetta and colleagues (*15*)) and an *ntnh*-like gene were identified in *Weissella oryzae* SG25, though no other genes associated with *bont* gene clusters were present (*7*). Also, a novel *bont*-like gene cluster (labelled BoNT/X) including *orfX*-like genes and a p47-like gene was recently identified on the chromosome of *C. botulinum* strain 111 (*8*). Mansfield and colleagues (*9*) recently identified distantly related lineages of *bont*-like sequences in *Chryseobacterium piperi*. These findings and the descriptions of horizontal gene transfer and recombination of botulinum neurotoxin genes and toxin gene clusters (*16*-*19*) suggest that the identification and diversity of novel botulinum-neurotoxin-like genes and gene clusters will expand with additional whole genome sequencing.

Here, we have identified a botulinum-neurotoxin-like gene cluster putatively located on a plasmid within the genome assembly of *Enterococcus* sp. 3G1_DIV0629. We bioinformatically characterized the *Enterococcus* sp. 3G1_DIV0629 genome as well as the gene and protein sequences associated with the *bont*-like gene cluster. This represents the first instance of BoNT-like sequences found in an *Enterococcus* sp. and suggests that homologous sequence to BoNTs may be more prevalent in the environment than previously appreciated.

## Methods

Known BoNT sequences were queried against the GenBank nr database with blastp (*20*, *21*). Hits were identified for *Enterococcus* sp. 3G1_DIV0629 (GcA_002141285.1). The genome assembly and sequencing data (SRR5645157,SRR5648109) for *Enterococcus* sp. 3G1_DIV0629 were downloaded from NCBI. To generate our own assembly, sequence data were assembled with the SPAdes assembler (*22*) and assemblies were annotated with Prokka (*23*). To test for potential laboratory constructs, contigs containing *bont*-like sequences were screened for vector contamination with VecScreen (https://www.ncbi.nlm.nih.gov/tools/vecscreen/). The contigs (nucleotide sequences) were compared to the NCBI nt database with blastn to gain insight into the origin of the contigs of interest (taxonomic origin and “genomic origin” – chromosome, plasmid or bacteriophage). Contigs were also compared to known plasmid sequences with progressiveMauve (*24*). Translated protein sequences of coding regions identified on contigs containing *bont*-like sequences were compared to the UniProt database (*25*) and the nr database with blastp to gain insight into the origin and function of proteins encoded on the contigs of interest.

Sequence data representing known BoNT serotypes and subtypes were downloaded from NCBI (Table 1). Genomic sequences containing botulinum neurotoxin genes were annotated with Prokka, and nucleotide and protein sequences of interest were extracted from Prokka output files. Gene cluster diagrams were generated with the R package genoPlotR (*27*). Aligned nucleotide sequences were compared with SimPlot (*28*) using the hamming distance option and filtering positions containing gaps in >75% of sequences to determine if the genes were the result of recombination. The diversity of the botulinum-neurotoxin-like sequences was also evaluated on the protein level as protein sequences are often more conserved than nucleotide sequences, which can be useful for comparing distantly related sequences. Botulinum neurotoxin protein sequences and BoNT-like protein sequences were aligned with MUSCLE, and a maximum likelihood phylogeny was inferred with IQ-TREE v1.5.5 (*26*) using the predicted best-fit model – VT+F+R4. The phylogeny was viewed with FigTree v1.4.3 (http://tree.bio.ed.ac.uk/software/figtree/). Protein sequences representing known BoNTs and newly identified BoNT-like sequences were compared with blastp. Protein sequences of novel and established BoNTs were aligned with MUSCLE (*29*) and the alignment was viewed in MEGA (*30*) to evaluate motifs of interest.

**Table 1.** Sequence data representing known botulinum neurotoxin serotypes and toxin subtypes

| Accession | Organism | Strain | BoNT | toxin_cluster_type |
| --- | --- | --- | --- | --- |
| ABDO02000001.1 | *C. botulinum* | NCTC2916 | A1 | orfX |
| AM412317.1 | *C. botulinum* | ATCC3502 | A1 | ha |
| CP006907.1 | *C. botulinum* | CDC297 | A1 | orfX |
| CP001581.1 | *C. botulinum* | Kyoto-F | A2 | orfX |
| CP000963.1 | *C. botulinum* | LochMaree | A3 | orfX |
| CP001081.1 | *C. botulinum* | 657 | A4 | orfX |
| FR773526.1 | *C. botulinum* | HO4402-65 | A5 | ha |
| FJ981696.1 | *C. botulinum* | CDC41370 | A6 | orfX |
| JQ954969.1 | *C. botulinum* | 2008-148 | A7 | orfX |
| KM233166.1 | *C. botulinum* | Chemnitz | A8 | orfX |
| CP000940.1 | *C. botulinum* | Okra | B1 | ha |
| AB855771.1 | *C. botulinum* | 111 | B2 | ha |
| LFQV01000032.1 | *C. botulinum* | CDC795 | B3 | ha |
| CP001057.1 | *C. botulinum* | Eklund17B | B4 | ha |
| CP001081.1 | *C. botulinum* | 657 | B5 | ha |
| BA000059.2 | *C. botulinum* | Osaka05 | B6 | ha |
| JQ354985.1 | *C. botulinum* | Bac-04-07755 | B7 | ha |
| JQ964806.1 | *C. botulinum* | Maehongson2010 | B8 | ha |
| ABDO02000001.1 | *C. botulinum* | NCTC2916 | (B) | ha |
| AP008983.1 | *C. botulinum* | c-st-Stockholm | C | ha |
| AB745667.1 | *C. botulinum* | 348 | CD | ha |
| AB012112.1 | *C. botulinum* | 1873-phage | D | ha |
| AB745668.1 | *C. botulinum* | OFD05 | DC | ha |
| ACSC01000002.1 | *C. botulinum* | Beluga | E1 | orfX |
| KF861920.1 | *C. botulinum* | FWKR11E1 | E10 | orfX |
| KF861879.1 | *C. botulinum* | SW280E | E11 | orfX |
| KF929215.1 | *C. botulinum* | 84-10 | E12 | orfX |
| EF028404.1 | *C. botulinum* | CDC5247 | E2 | orfX |
| CP001078.1 | *C. botulinum* | AlaskaE43 | E3 | orfX |
| ACOM01000005.1 | *C. botulinum* | BL5262 | E4 | orfX |
| AB037704.1 | *C. botulinum* | LCL155 | E5 | orfX |
| AM695752.1 | *C. botulinum* | K35 | E6 | orfX |
| JN695729.1 | *C. botulinum* | IBCA97-0192 | E7 | orfX |
| JN695730.1 | *C. botulinum* | Bac-02-06430 | E8 | orfX |
| ALYJ01000070.1 | *C. botulinum* | CDC66177 | E9 | orfX |
| CP000728.1 | *C. botulinum* | Langeland | F1 | orfX |
| ABDP01000023.1 | *C. botulinum* | Bf | F2 | orfX |
| GU213227.1 | *C. botulinum* | VPI4257-F160 | F3 | orfX |
| AOSX01000018.1 | *C. botulinum* | Af84 | F4 | orfX |
| AOSX01000021.1 | *C. botulinum* | Af84 | F5 | orfX |
| CP006903.1 | *C. botulinum* | Eklund202F | F6 | orfX |
| CP006905.1 | *C. botulinum* | Sullivan | F7 | orfX |
| AUZC01000009.1 | *C. botulinum* | I357 | F8 | orfX |
| AYSO01000020.1 | *C. botulinum* | CDC2741 | G | ha |
| JSCF01000006.1 | *C. botulinum* | CFSAN024410 | HA | orfX |
| AF528097.1 | *C. botulinum* | E88 | tetanus | NA |
| AP014696.1 | *C. botulinum* | 111 | bont-like | orfX |
| BAWR01000013.1 | *Weissella oryzae* | SG25 | bont-like | NA |
| NGLI01000004.1 | *Enterococcus sp.* | 3G1-DIV0629 | bont-like | orfX |

To determine the phylogenetic relationship of strain 3G1_DIV0629 to other *Enterococcus* spp., the publicly available assembly and the in-house assembly (SPAdes) were compared to RefSeq genomes using Mash v2.0 (*31*) and the presketched RefSeq archive (https://gembox.cbcb.umd.edu/mash/refseq.genomes%2Bplasmid.k21s1000.msh). The contig containing *bont*-like toxin cluster sequences from both assemblies were also compared to the pre-sketched RefSeq and plasmid databases. Whole genome SNP analysis was used to further investigate the relationship of *Enterococcus* sp. 3G1_DIV0629 with other *Enterococcus faecium* isolates. *E. faecium* genome assemblies were downloaded from NCBI and SNPs were called from NUCmer (*32*, *33*) alignments to a reference genome (GCA_000174395.2) within NASP (*34*). SNPs identified in duplicated regions of the reference genome (with NUCmer self-alignments) were filtered from downstream analyses. A maximum likelihood phylogeny was generated from the concatenated SNP data (114904 positions) with IQ-TREE v1.5.5 (*26*) using the predicted best-fit model – TVM+F+AsC+R3.

Supplemental data files including the in-house genome assembly and BoNT protein sequences evaluated in this study are available on github – https://github.com/chawillia/bont-like-sequences_2017.git.

## Results

Blastp searches of BoNT sequences against publicly available databases revealed a BoNT-like protein from the recently whole genome sequenced *Enterococcus* sp. 3G1_DIV0629. The *bont*-like gene sequence was associated with a gene cluster that included an *ntnh*-like gene, *orfX*-like genes and a p47-like gene, which is similar to known botulinum neurotoxin gene clusters (*10*) (Figure 1). While the order and orientation of the genes in the *Enterococcus* sp. 3G1_DIV0629 cluster are similar to known *orfX*+ botulinum neurotoxin gene clusters, only two of the three *orfX* genes (orf-X2 and orf-X3) are present in the *bont*-like gene cluster of *Enterococcus* sp. 3G1_DIV0629, and these genes are in the opposite orientation of *orfX* genes in known botulinum neurotoxin gene clusters. This orientation of the *orfX* genes is similar to a recently identified *bont*-like gene cluster on the chromosome of *C. botulinum strain* 111 (*8*) (Figure 1). A SimPlot comparing aligned nucleotide sequences (Figure 2) indicates a consistent level of identity along the length of the *bont*-like gene sequence when compared to known botulinum neurotoxin genes, which suggests that the gene is not the result of a novel recombination of multiple serotypes. Zhang and colleagues (*8*) made similar observations when evaluating the *bont*-like sequence in *C. botulinum* strain 111.

**Figure 1.**
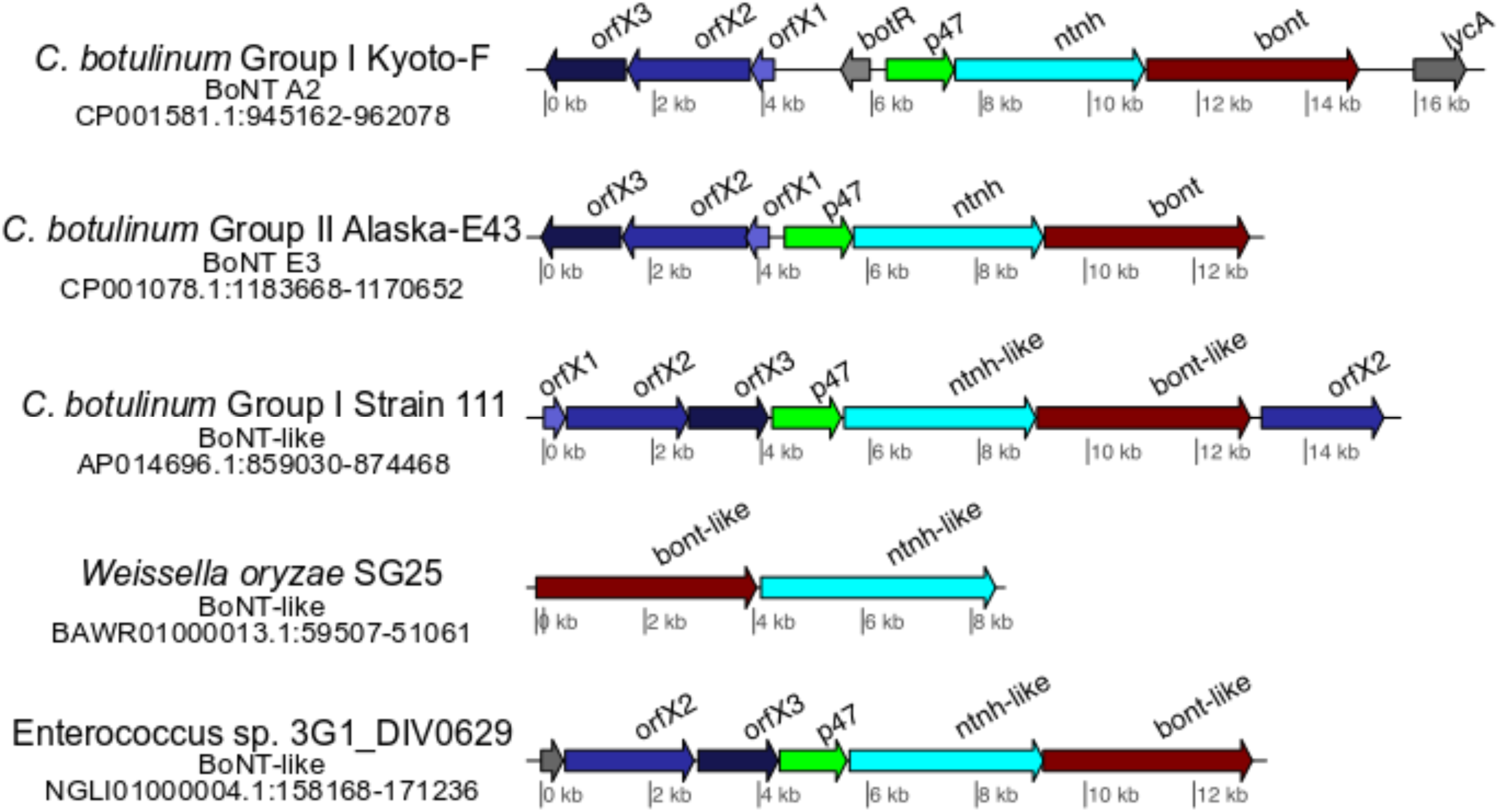
Structure of botulinum neurotoxin gene clusters and *bont*-like gene clusters. Toxin gene clusters of *orfX*+ strains Kyoto-F and Alaska-E43 are displayed for reference. Searching publicly available data has resulted in the identification of *bont*-like toxin gene clusters in *C. botulinum* Strain 111 (*8*), *W. oryzae* SG25 (*7*) and *Enterococcus* sp. 3G1_DIV0629 (this paper). The *bont*-like gene cluster in *Enterococcus* sp. 3G1_DIV0629 contains an *ntnh*-like and *bont*-like gene similar to known botulinum neurotoxin gene clusters. *orfX2* and *orfX3*-like genes are in a different orientation than botulinum neurotoxin gene clusters in strains known to cause botulism (e.g. Kyoto-F and Alaska E43). The *bont*-like toxin gene cluster arrangement in *Enterococcus* sp. 3G1_DIV0629 is similar to the arrangement of the *bont*-like gene cluster recently identified in *C. botulinum* Strain 111 (*8*).

**Figure 2.**
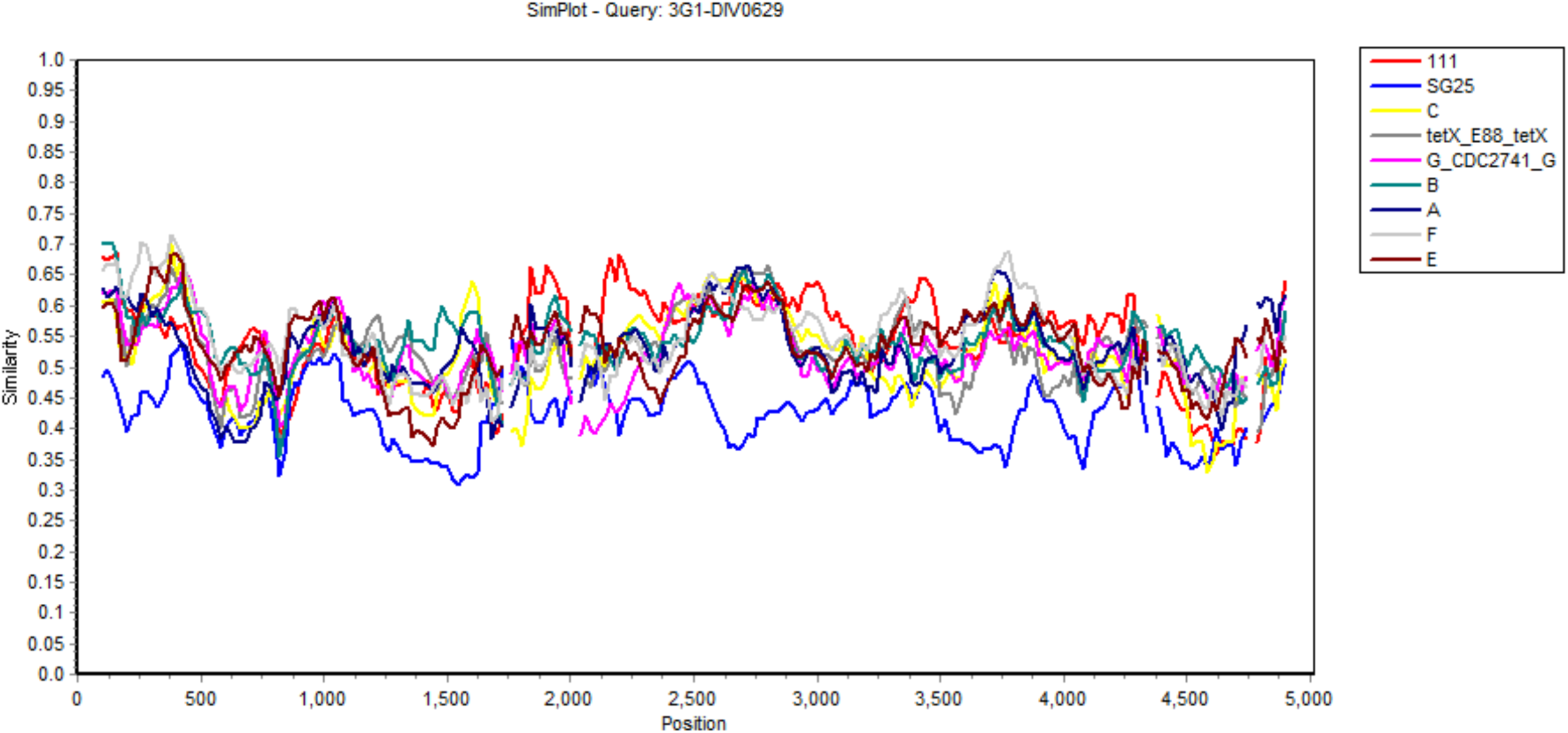
A SimPlot comparing an alignment of known botulinum neurotoxin gene sequences (MUSCLE alignment – 5014 positions) to the *bont*-like gene sequence from *Enterococcus* sp. 3G1_DIV0629 (reference sequence for comparisons). SimPlot was run using the hamming distance option and filtering positions containing gaps in >75% of sequences. The *bont*-like gene (*bont*/X) from *C. botulinum* strain 111 is displayed in red; this sequence is the most closely related to the *Enterococcus* sp. 3G1_DIV0629 sequence of the evaluated sequences according to the BoNT phylogeny (Figure 3) and blastp results (Table 2) when comparing BoNT protein sequences.

**Table 2.** Blastp comparisons of botulinum neurotoxins and associated protein

|  |  |  |  |  | 3G1-DIV0629 sequences queried against references (%ID, coverage) |  |  |  |  |
| --- | --- | --- | --- | --- | --- | --- | --- | --- | --- |
| Accession | Organism | Strain | BoNT | toxin_cluster_type | BoNT | NTNH | P47 | orfx3 | orfx2 |
| AM412317.1 | *C. botulinum* | ATCC3502 | A1 | ha | 29.12,99 | 27.42,97 |  |  |  |
| CP006907.1 | *C. botulinum* | CDC297 | A1 | orfX | 29.28,99 | 26.52,97 | 30,61 | 36.62,99 | 23.84,99 |
| ABDO02000001.1 | *C. botulinum* | NCTC2916 | A1 | orfX | 29.21,99 | 26.68,97 | 30.26,61 | 36.62,99 | 23.84,99 |
| CP001581.1 | *C. botulinum* | Kyoto-F | A2 | orfX | 29.12,99 | 26.74,97 | 30,61 | 36.62,99 | 23.41,99 |
| CP000963.1 | *C. botulinum* | LochMaree | A3 | orfX | 28.93,99 | 27.36,97 | 30.37,61 | 37.02,99 | 23.97,95 |
| CP001081.1 | *C. botulinum* | 657 | A4 | orfX | 29.3,97 | 26.88,96 | 26.54,82 | 36.2,99 | 25.03,99 |
| FR773526.1 | *C. botulinum* | HO4402-65 | A5 | ha | 29.12,99 | 26.98,97 |  |  |  |
| FJ981696.1 | *C. botulinum* | CDC41370 | A6 | orfX | 29.2,99 | 26.74,97 | 30,61 |  | 23.65,99 |
| JQ954969.1 | *C. botulinum* | 2008-148 | A7 | orfX | 28.77,99 |  |  |  |  |
| KM233166.1 | *C. botulinum* | Chemnitz | A8 | orfX | 30.04,97 | 26.57,96 | 30.63,70 | 36.2,99 | 23.53,99 |
| CP000940.1 | *C. botulinum* | Okra | B1 | ha | 29.27,99 | 27.09,98 |  |  |  |
| AB855771.1 | *C. botulinum* | 111 | B2 | ha | 29.14,99 | 26.93,98 |  |  |  |
| LFQV01000032.1 | *C. botulinum* | CDC795 | B3 | ha | 29.08,99 | 27.17,98 |  |  |  |
| CP001057.1 | *C. botulinum* | Eklund17B | B4 | ha | 29.19,99 | 27.06,97 |  |  |  |
| CP001081.1 | *C. botulinum* | 657 | B5 | ha | 29.4,99 | 27.09,98 |  |  |  |
| BA000059.2 | *C. botulinum* | Osaka05 | B6 | ha | 29.12,99 | 26.93,98 |  |  |  |
| JQ354985.1 | *C. botulinum* | Bac-04-07755 | B7 | ha | 29.22,99 |  |  |  |  |
| JQ964806.1 | *C. botulinum* | Maehongson2010 | B8 | ha | 29.22,99 | 27.01,98 |  |  |  |
| ABDO02000001.1 | *C. botulinum* | NCTC2916 | (B) | ha |  | 27.3,98 |  |  |  |
| AP008983.1 | *C. botulinum* | c-st-Stockholm | C | ha | 29.56,86 | 25.55,97 |  |  |  |
| AB745667.1 | *C. botulinum* | 348 | CD | ha | 29.38,90 | 26.1,97 |  |  |  |
| AB012112.1 | *C. botulinum* | 1873-phage | D | ha | 29.76,91 | 25.63,97 |  |  |  |
| AB745668.1 | *C. botulinum* | OFD05 | DC | ha | 29.13,86 | 25.96,97 |  |  |  |
| ACSC01000002.1 | *C. botulinum* | Beluga | E1 | orfX | 28.5,99 | 26.93,94 | 29.96,61 | 37.04,99 | 26.08,94 |
| KF861920.1 | *C. botulinum* | FWKR11E1 | E10 | orfX | 28.84,99 |  |  |  |  |
| KF861879.1 | *C. botulinum* | SW280E | E11 | orfX | 28.58,99 |  |  |  |  |
| KF929215.1 | *C. botulinum* | 84-10 | E12 | orfX | 28.38,99 |  |  |  |  |
| EF028404.1 | *C. botulinum* | CDC5247 | E2 | orfX | 28.49,99 |  |  |  |  |
| CP001078.1 | *C. botulinum* | AlaskaE43 | E3 | orfX | 28.37,99 | 26.93,94 | 29.96,61 | 37.04,99 | 26.17,94 |
| ACOM01000005.1 | *C. botulinum* | BL5262 | E4 | orfX | 28.69,99 | 26.33,96 | 30.34,61 | 37.04,99 | 25.78,94 |
| AB037704.1 | *C. botulinum* | LCL155 | E5 | orfX | 28.25,99 |  |  |  |  |
| AM695752.1 | *C. botulinum* | K35 | E6 | orfX | 28.89,99 | 26.78,96 | 29.96,61 |  | 26.04,94 |
| JN695729.1 | *C. botulinum* | IBCA97-0192 | E7 | orfX | 28.74,99 |  |  |  |  |
| JN695730.1 | *C. botulinum* | Bac-02-06430 | E8 | orfX | 28.76,99 |  |  |  |  |
| ALYJ01000070.1 | *C. botulinum* | CDC66177 | E9 | orfX | 28.74,99 | 27,96 | 30.34,61 | 36.69,99 | 25.03,94 |
| CP000728.1 | *C. botulinum* | Langeland | F1 | orfX | 28.86,99 | 26.38,99 | 30.63,70 | 36.2,99 | 23.53,99 |
| ABDP01000023.1 | *C. botulinum* | Bf | F2 | orfX | 29.18,99 | 26.93,94 | 26.54,82 | 36.85,99 | 22.51,99 |
| GU213227.1 | *C. botulinum* | VPI4257-F160 | F3 | orfX | 29.19,99 |  |  |  |  |
| AOSX01000018.1 | *C. botulinum* | Af84 | F4 | orfX | 29.13,99 | 26.54,94 | 30.37,61 | 36.09,99 | 26.36,94 |
| AOSX01000021.1 | *C. botulinum* | Af84 | F5 | orfX | 29.31,99 | 26.91,98 | 30,61 | 37.02,99 | 23.97,95 |
| CP006903.1 | *C. botulinum* | Eklund202F | F6 | orfX | 29.59,99 | 26.61,98 | 30.71,61 | 36.82,99 | 22.29,95 |
| CP006905.1 | *C. botulinum* | Sullivan | F7 | orfX | 27.76,99 | 26.1,99 | 32.21,61 | 35.7,99 | 26.88,95 |
| AUZC01000009.1 | *C. botulinum* | I357 | F8 | orfX | 29.14,99 | 26.96,98 | 29.26,61 | 36.87,99 | 23.96,94 |
| JSCF01000006.1 | *C. botulinum* | CFSAN024410 | HA | orfX | 28.42,99 | 26.71,99 | 29.89,61 | 37.22,99 | 23.38,99 |
| AYSO01000020.1 | *C. botulinum* | CDC2741 | G | ha | 28.79,99 | 25.78,99 |  |  |  |
| AF528097.1 | *C. botulinum* | E88 | tetanus | NA | 28.89,99 |  |  |  |  |
| AP014696.1 | *C. botulinum* | 111 | bont-like | orfX | 38.54,99 | 29.89,99 | 33.09,61 | 39.16,99 | 27.57,98/25.72,100 |
| BAWR01000013.1 | *Weissella oryzae* | SG25 | bont-like | NA | 26.37,49 | 23.21,37 |  |  |  |
| NGLI01000004.1 | *Enterococcus sp.* | 3G1-DIV0629 | bont-like | orfX |  |  |  |  |  |

Sequence data representing each of the botulinum neurotoxin serotypes and subtypes were downloaded from NCBI (Table 1) and predicted protein sequences were compared. A phylogeny of botulinum neurotoxin protein sequences (Figure 3), including the BoNT-like sequence described here, indicates that the BoNT-like protein predicted from the assembly for *Enterococcus* sp. 3G1_DIV0629 is not closely related to known BoNTs that cause most botulism cases in humans (serotypes A and B). The BoNT-like sequence is most closely related to a BoNT-like protein sequence recently identified on the *C. botulinum* Group I Strain 111 chromosome (*8*). The predicted protein sequences of the *bont*-like toxin cluster genes in *Enterococcus* sp. 3G1_DIV0629 were also queried against known botulinum toxins and associated proteins with blastp (Table 2). The BoNT-like protein sequence of *Enterococcus* sp. 3G1_DIV0629 generally shares approximately 29% (median=29.12%, median coverage of 97% - Table 2) identity with known BoNT proteins. However, the BoNT-like sequence from *Enterococcus* sp. 3G1_DIV0629 shares 38.54% identity with the BoNT-like sequence from *C. botulinum* strain 111. As observed with the BoNT-like protein sequences in *W. oryzae* SG25 and *C. botulinum* strain 111 (*7*, *8*), the HExxH metallopeptidase motif (HELCH in *Enterococcus* sp. 3G1_DIV0629) is conserved in the predicted BoNT-like protein sequence of *Enterococcus* sp. 3G1_DIV0629 (Figure 4A). Additionally, the SxWY binding motif is conserved in the BoNT-like protein sequence of *Enterococcus* sp. 3G1_DIV0629 (Figure 4B) which suggests the protein could potentially interact with neuronal cells. The NTNH-like, P47-like and ORFX-like protein sequences also show relatively low sequence identity to protein sequences associated with known BoNTs (Table 2).

**Figure 3.**
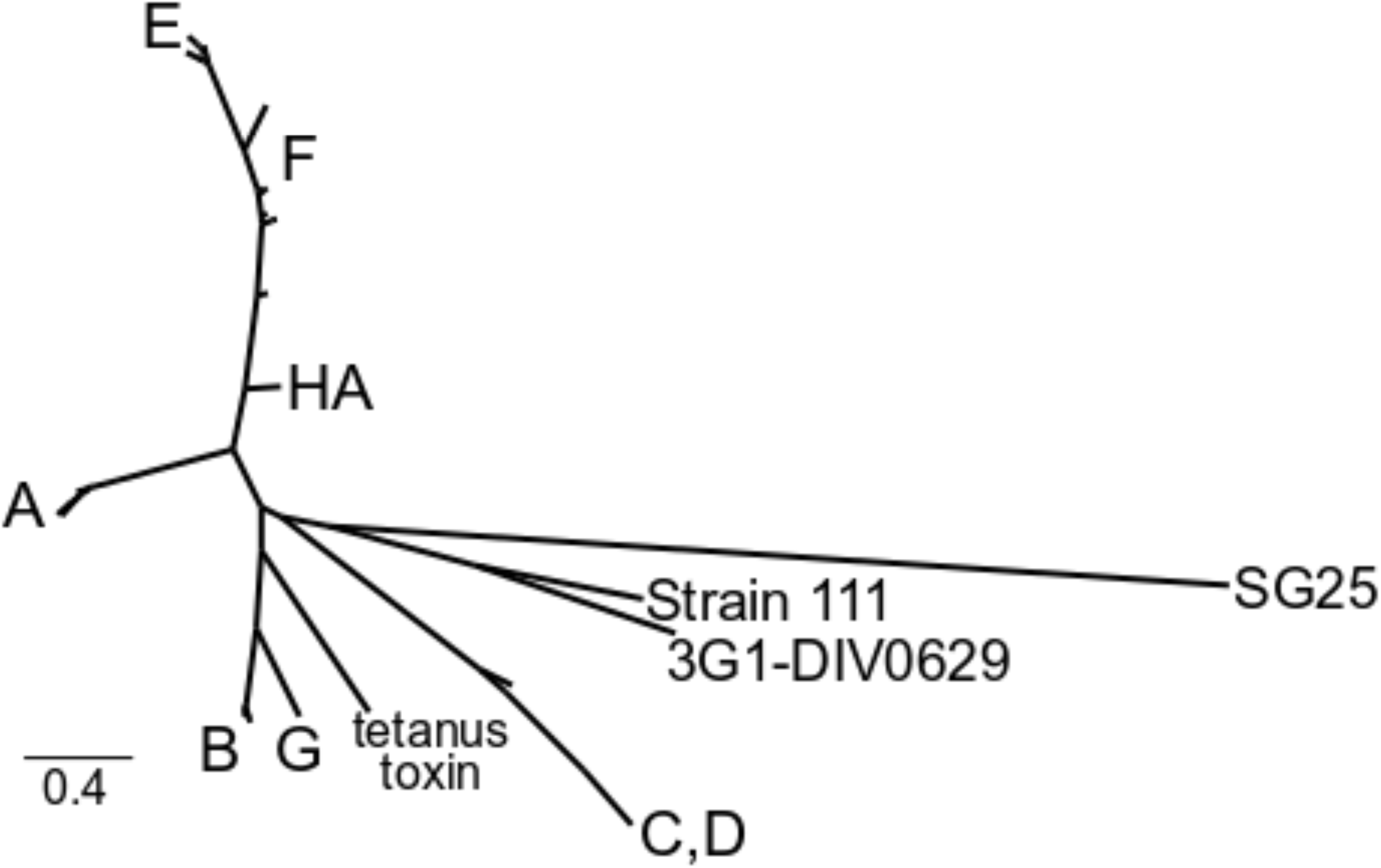
Maximum likelihood phylogeny of botulinum neurotoxin proteins and recently identified BoNT-like protein sequences. Forty-eight protein sequences representing known botulinum neurotoxins and recently identified BoNT-like sequences were aligned with MUSCLE and a phylogeny was inferred with IQ-TREE.

**Figure 4.**
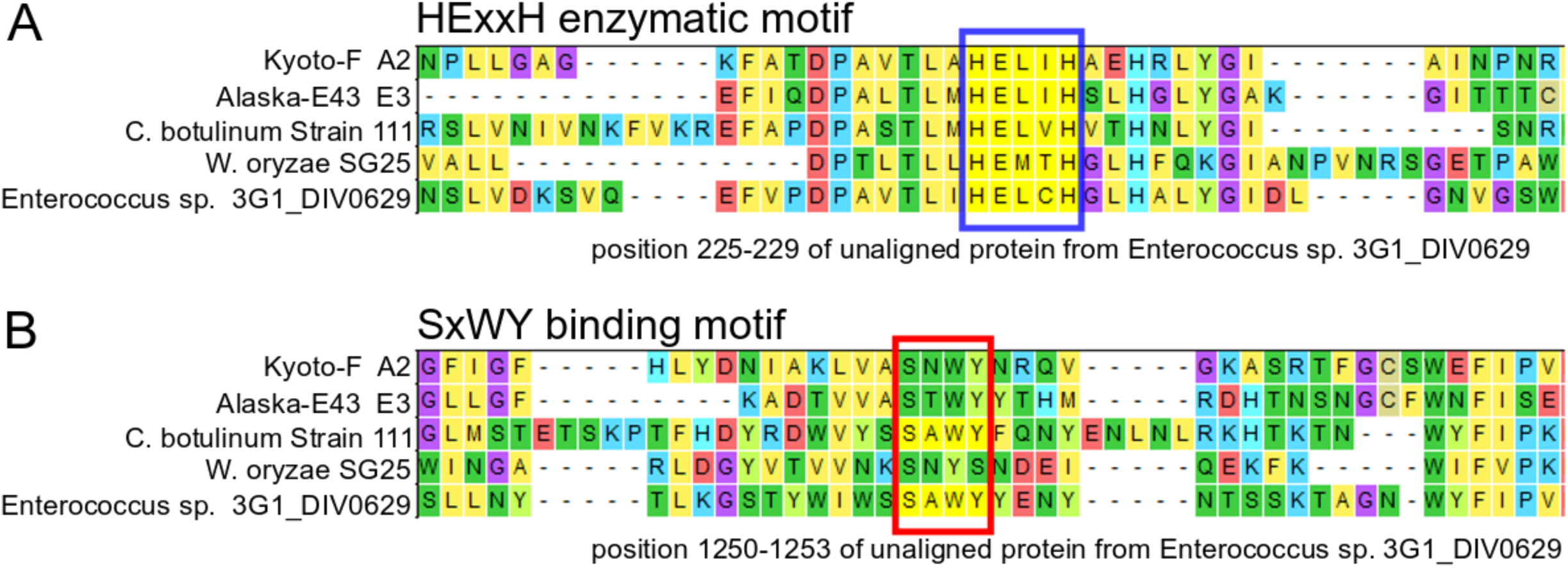
Muscle alignment of BoNT sequences viewed with Mega. (A) The HExxH protease motif that is conserved among known BoNTs and the BoNT-like protein reported here is identified by the blue box. (B) The red box identifies the SxWY motif that plays a role in ganglioside binding. This motif is conserved in known BoNTs and its presence in the BoNT-like protein identified in *Enterococcus* sp. 3G1_DIV0629 suggests that the protein could potentially interact with neuronal cells.

The genome assembly for *Enterococcus* sp. 3G1_DIV0629 was investigated to characterize the contig containing the *bont*-like gene cluster and to determine the relationship of the strain to other *Enterococcus* spp. Contigs containing *bont*-like sequences, NGLI01000004.1 and NODE_32 from an in-house assembly (SPAdes), show no signs of vector contamination when screened with VecScreen against the UniVec database. Several lines of evidence suggest the *bont*-like gene cluster has been inserted into the *Enterococcus* genomic background and is putatively located on a plasmid. The GC content of the contigs containing the *bont*-like sequences is slightly lower than the GC content of the entire genome assembly (Table 3). Comparisons of bacterial chromosomes with plasmids and insertion sequences have shown that the GC content of the chromosome is often higher than the GC content of the associated plasmid or insertion sequence (*35*, *36*). Comparisons of the contigs containing the *bont*-like sequences to reference genomes and plasmids with Mash distances suggest that the contigs are most closely related to *Enterococcus* spp. plasmid sequences. Additionally, blastn comparisons of the entire contigs as well as the regions adjacent to the *bont*-like gene cluster share identity with sequences associated with *Enterococcus* spp. plasmids. Contigs from an in-house genome assembly for *Enterococcus* sp. 3G1_DIV0629 were compared to *E. faecium* plasmids with progressiveMauve (Figure 5). The regions surrounding the *bont*-like gene cluster share homology with known *Enterococcus* plasmid sequences. However, the genomic location of the *bont*-like gene cluster is only putative due to the draft nature of the genome assembly. Coding region sequences in the regions adjacent to the *bont*-like gene cluster are annotated as transposases, which could explain how the *bont*-like genes were inserted into the *Enterococcus* genomic background.

**Figure 5.**
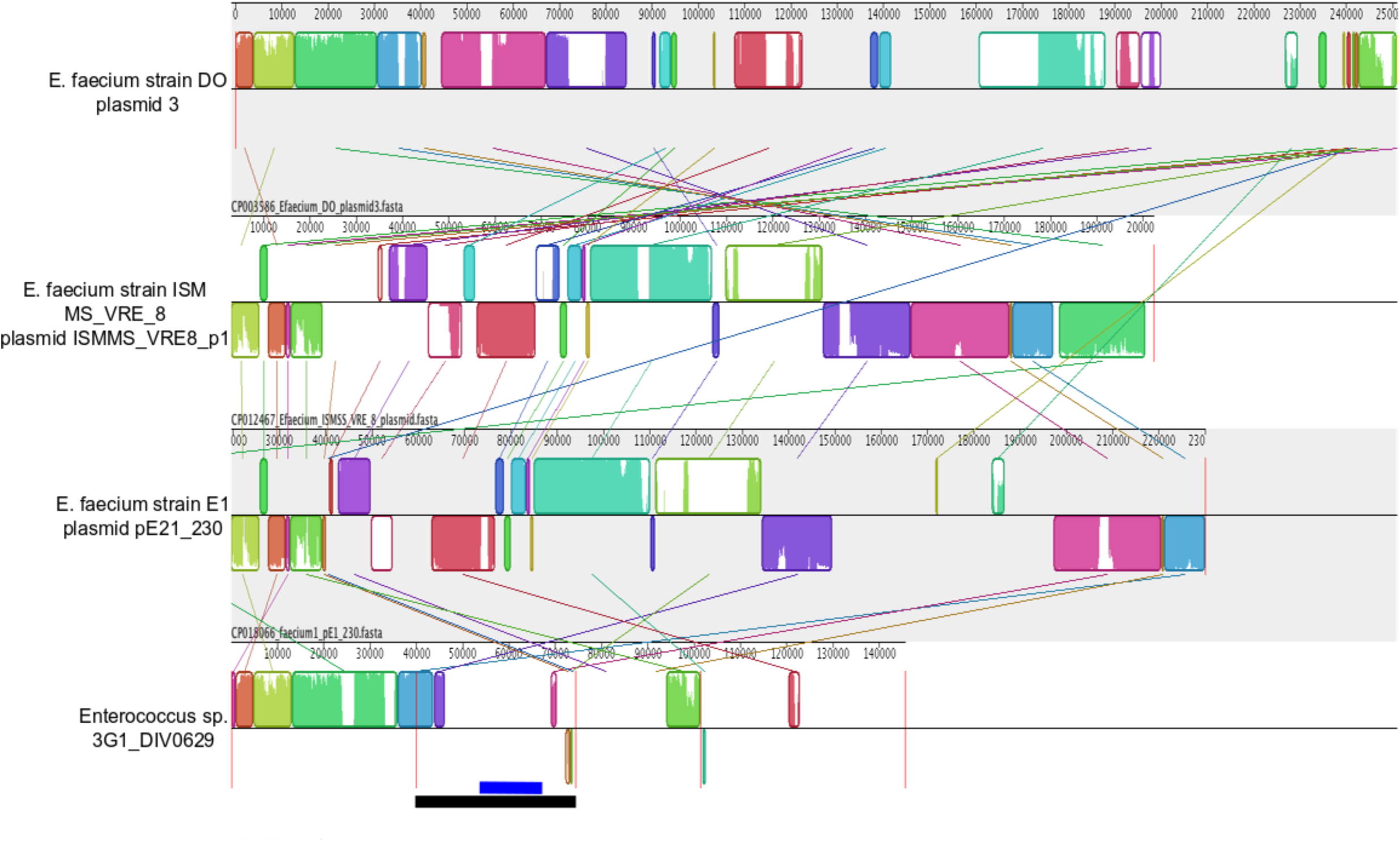
Comparison of *E. faecium* plasmid sequences and contigs from an in-house assembly for *Enterococcus* sp. 3G1_DIV0629 with progressiveMauve. Four contigs from the in-house assembly (NODE_26, NODE_32, NODE_42, NODE_24) with BLAST hits to *E. faecium* plasmid sequences were arbitrarily concatenated into a single fasta file for the comparison. NODE_32 contains the *bont*-like gene cluster for *Enterococcus* sp. 3G1_DIV0629 – the black line at the bottom of the figure indicates this contig, and the blue line indicates the approximate location of the *bont*-like gene cluster. The regions surrounding the *bont*-like gene cluster share homology with known *E. faecium* plasmids, which suggests the *bont*-like gene cluster is present on a plasmid. At the edges of the regions that share homology with known plasmids, there are coding regions annotated as transposases.

**Table 3.** GC content of genome assemblies and contigs of interest

|  | GC content (%) |  |
| --- | --- | --- |
| Genome Assembly | Entire Assembly | *bont*-like Sequence Contig |
| GCA_002141285.1 | 37.71 | 31.85 |
| SPAdes in-house assembly | 37.66 | 31.82 |

The draft genome assembly of *Enterococcus* sp. 3G1_DIV0629 was compared to publicly available genomic data to determine how this strain is related to other *Enterococcus* spp. Comparisons of the genome assembly against RefSeq genomes with Mash indicate that top hits for the *Enterococcus* sp. 3G1_DIV0629 assembly are members of *Enterococcus faecium.* Thus, *Enterococcus* sp. 3G1_DIV0629 was compared to *E. faecium* genomes with a core genome SNP analysis (Figure 6). The most closely related genomes to *Enterococcus* sp. 3G1_DIV0629 are *E. faecium* T110 (GCA_000737555.1) and *E. faecium* L-X (GCA_000787065.1), both of which are identified as probiotic strains.

**Figure 6.**
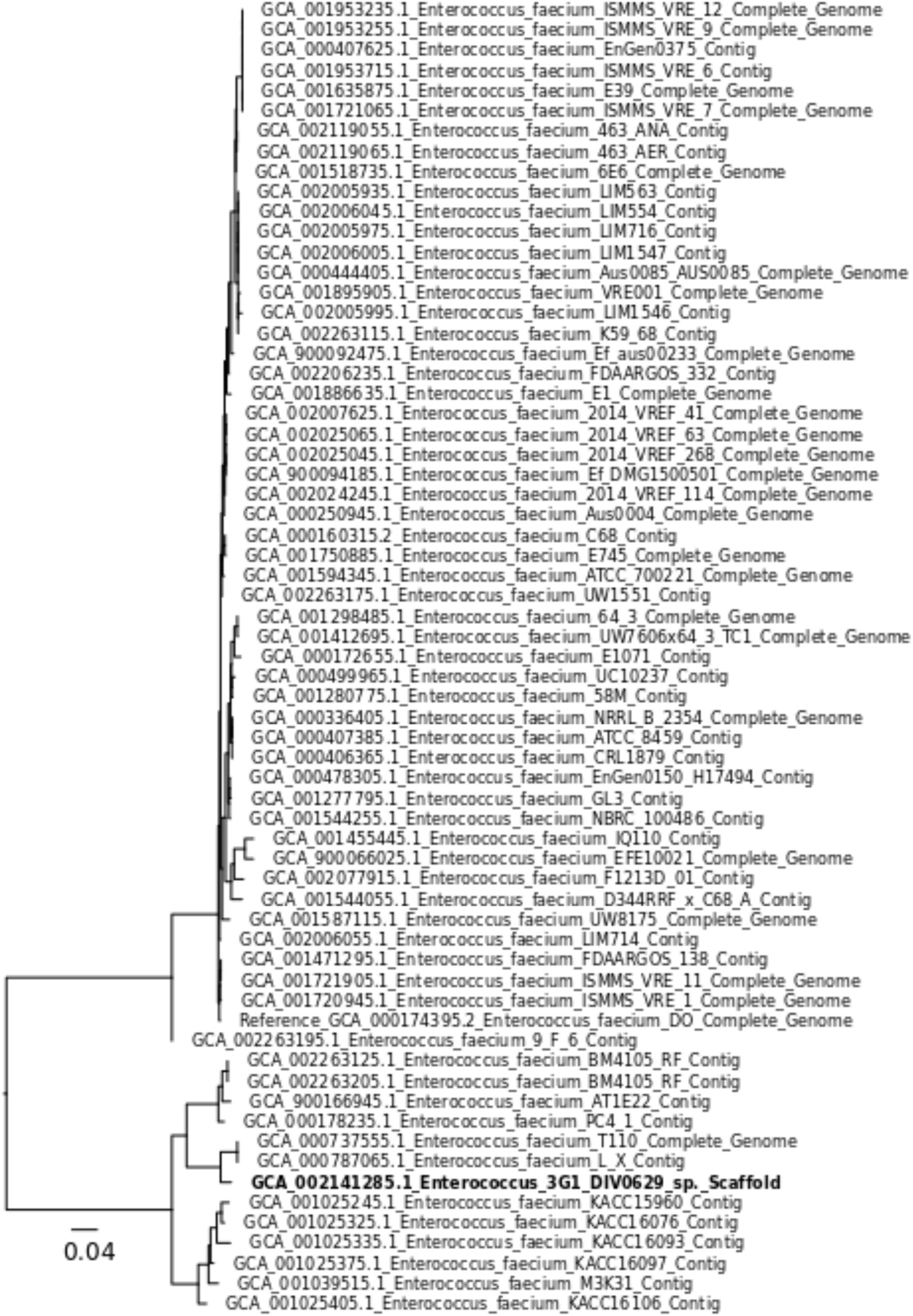
Maximum likelihood phylogeny of concatenated core genome SNPs. *Enterococcus* sp. 3G1_DIV0629 is indicated with bold text and is closely related to *E. faecium* isolates identified as probiotic strains.

## Discussion

Exploring the diversity of BoNTs is important for the development of therapeutics and antitoxins and also to understand the function, evolution, and pathogenesis of botulinum neurotoxins. Here we describe a novel, *bont*-like gene cluster that was identified in a draft genome assembly for *Enterococcus* sp. 3G1_DIV0629 by querying publicly available genomic databases. A similar data mining approach was recently used to identify a novel botulinum-neurotoxin-like gene cluster in a Group I *C. botulinum* genome, strain 111 (8). Metadata associated with the *Enterococcus* sp. 3G1_DIV0629 genome assembly indicate the bacterium was isolated from a bovine fecal sample in South Carolina, USA. The isolate is closely related to members of *E. faecium. E. faecium* strains are Gram-positive, non-endospore forming facultative anaerobes that are members of the Firmicutes and commonly inhabit the digestive tracts of mammals (*37*, *38*). *E. faecium* has been associated with antimicrobial resistance and a variety of human infections (*39*-*41*). Interestingly, antagonistic interactions between *E. faecium* and *C. botulinum* have been reported. *E. faecium* has been described to inhibit the growth of *C. botulinum* strains and inhibit BoNT production (*42*-*44*). The impact of ruminal microbial communities, particularly *Enterococcus* spp., on bovine botulism has been an area of on-going research (*45*-*48*).

This report is the first instance of identifying botulinum-neurotoxin-like sequences in a member of the *Enterococcus.* The *bont*-like gene cluster identified in *Enterococcus* sp. 3G1_DIV0629 is putatively located on a plasmid sequence, and the regions surrounding the *bont*-like gene cluster are putatively of *Enteroccocus* origin. Coding region sequences in the regions adjacent to the bont-like gene cluster are annotated as transposases. Horizontal gene transfer of botulinum neurotoxin genes via association with transposases has been described (*17*, *18*). One explanation for the presence of the *bont*-like gene cluster in *Enterococcus* sp. 3G1_DIV0629 is that the gene cluster was inserted from an unknown source into the *Enterococcus* genomic background (putatively on a plasmid) long ago and has since diverged from known botulinum neurotoxin genes. Alternatively, this gene cluster could be the result of a more recent insertion event from a divergent source (divergent from known BoNTs). The toxin-like gene cluster sequence could provide some unknown ecological advantage for survival in the environment or in a host. Similar *bont*-like sequences may be present in additional organisms that have yet to be sampled as they are difficult to culture or are not pertinent to studies regarding human health, which have driven a great deal of genome sequencing efforts.

The botulinum-neurotoxin-like gene cluster in *Enterococcus* sp. 3G1_DIV0629 was identified using bioinformatic search tools and publicly available genomic sequence data. The gene cluster is similar to known botulinum neurotoxin gene clusters that contain *orfX*+ genes. The HExxH protease motif and the SxWY binding motif that are conserved in known BoNTs are present in the predicted BoNT protein sequence. Importantly, it is currently unknown if the BoNT-like protein described here is capable of targeting neuronal cells resulting in botulism or if the BoNT-like protein and associated proteins are even expressed by the bacterium; there is likely a fitness cost with maintaining the *bont*-like gene cluster, which suggests that this region has an ecological function that has yet to be identified. No known cases of botulism have been attributed to *Enterococcus* spp. Thus, it seems unlikely that these microbes are expressing proteins that cause botulism in humans; additional experiments would be required to test this hypothesis. Bioinformatic studies such as this coupled with laboratory experiments can inform our understanding of the diversity and evolution of Clostridial toxins.

## Acknowledgements

Opinions, interpretations, conclusions and recommendations are those of the authors and not necessarily endorsed by the U.S. Army.

